# COSMIC-dFBA: A novel multi-scale hybrid framework for bioprocess modeling

**DOI:** 10.1101/2023.09.13.557646

**Authors:** Saratram Gopalakrishnan, William Johnson, Miguel A. Valderrama-Gomez, Elcin Icten, Jasmine Tat, Michael Ingram, Coral Fung Shek, Pik K. Chan, Fabrice Schlegel, Pablo Rolandi, Cleo Kontoravdi, Nathan Lewis

## Abstract

Metabolism governs cell performance in biomanufacturing, as it fuels growth and productivity. However, even in well-controlled culture systems, metabolism is dynamic, with shifting objectives and resources, thus limiting the predictive capability of mechanistic models for process design and optimization. Here, we present Cellular Objectives and State Modulation In bioreaCtors (COSMIC)-dFBA, a hybrid multi-scale modeling paradigm that accurately predicts cell density, antibody titer, and bioreactor metabolite concentration profiles. Using machine-learning, COSMIC-dFBA decomposes the instantaneous metabolite uptake and secretion rates in a bioreactor into weighted contributions from each cell state (growth or antibody-producing state) and integrates these with a genome-scale metabolic model. A major strength of COSMIC-dFBA is that it can be parameterized with only metabolite concentrations from spent media, although constraining the metabolic model with other omics data can further improve its capabilities. Using COSMIC-dFBA, we can predict the final cell density and antibody titer to within 10% of the measured data, and compared to a standard dFBA model, we found the framework showed a 90% and 72% improvement in cell density and antibody titer prediction, respectively. Thus, we demonstrate our hybrid modeling framework effectively captures cellular metabolism and expands the applicability of dFBA to model the dynamic conditions in a bioreactor.

## 1. Introduction

Maximizing recombinant protein titer in a pharmaceutical bioprocess can be facilitated by optimizing nutrient feeding. Optimal conditions are commonly identified using time and resource-intensive design of experiments (DOE) strategies (Kasemiire et al., 2021). Models built on process data can help predict the trajectory of cellular states and control the process environment (Sidoli et al., 2004). Predictive models have previously leveraged empirical Monod-based equations to compute growth rates based on the extracellular concentrations of limiting nutrients (Ben Yahia et al., 2021; Galleguillos et al., 2017). The uptake and secretion rates for non-limiting nutrients are described by their relative uptake/secretion rates and/or kinetic rate laws defined by concentrations (López-Meza et al., 2016). However, nutrient depletion and toxic metabolite accumulation leads to metabolic shifts that cause uptake and secretion rates relative to the limiting nutrient to change during the bioprocess (Sunley et al., 2008; Templeton et al., 2013). This limitation motivates the inclusion of descriptive and mechanistic models of cellular metabolism in dynamic bioreactor models.

Genome-scale metabolic models are comprehensive collections of all metabolic pathways for an organism and are valuable for predicting product yields when nutrient uptake rates are specified. Metabolic flux through the entire network can be predicted using constraint-based modeling, such as flux balance analysis (Orth et al., 2010), which assumes that resource allocation in a cell aims to fulfill specific cellular objectives. This capability is leveraged for dynamic flux balance analysis (dFBA) (Mahadevan et al., 2002) and uses bioreactor substrate concentrations to determine nutrient uptake by the metabolic model. Fluxes are then predicted with the metabolic model to update metabolite concentrations in the bioreactor. Overall, this framework embeds the FBA problem within a system of ODEs to predict metabolic and cellular dynamics in the reactor.

While dFBA is structurally simple, it has three disadvantages that limit its application to mammalian bioprocessing. First, cellular metabolism is dynamic and therefore, the metabolic model must be tailored to be consistent with the extracellular environment. Otherwise, the full genome-scale model over-predicts intracellular fluxes as it affords the use of conditionally inactivated pathways (Jerby et al., 2010). Second, changes in extracellular environments cause cells to change the abundance of transporter proteins, which further changes kinetic parameters governing nutrient uptake rates (Laakso et al., 2011). Third, cells exhibit metabolic shifts arising from metabolite accumulation, such as lactate, wherein lactate production switches to lactate consumption during the bioprocess (Torres et al., 2018). This is frequently seen in fed-batch cultures with CHO cells and must be conditionally integrated into existing bioprocess models (Nolan and Lee, 2011).

Capturing metabolic shifts requires us to first characterize them. Some algorithms rely on visual inspection (Dean and Reddy, 2013) or piecewise linear regression (Ben Yahia et al., 2017) to identify different process phases. However, these methods suffer from the drawback that the model may reflect a single dataset or growth condition. Thus, they may not generalize to other conditions prevalent in the bioreactor or states of a bioprocess. Finally, predicting product fluxes requires us to know a cell’s objectives for a given cellular state. Objective functions, such as growth rate maximization, can be reliably applied to quantify metabolism in prokaryotes; however, these objectives have limited relevance to mammalian cells, since they only partially characterize the growth phase (Savinell and Palsson, 1992). To model the non-growing states, alternative objective functions must be explored (Garcia Sanchez and Torres Saez, 2014). More recently, parsimonious nutrient uptake was proposed as an objective (Chen et al., 2019), but it does not capture the variation in amino acid allocation towards different recombinant proteins.

Therefore, there is a need for a comprehensive framework that correctly and models the biological characteristics of the cells in the bioreactor with high fidelity by addressing the changes in cell states arising from constantly changing conditions in an industrial bioprocess.

Here we present Cellular Objectives and State Modulation In bioreaCtors (COSMIC)-dFBA, a multi-scale modeling framework for predicting concentration profiles of glucose, metabolic byproducts, antibody, amino acids, and cell density in a perfusion bioprocess (Figure 1). As with standard dFBA, COSMIC-dFBA predicts concentration profiles of metabolites by solving a system of ODE equations in which the uptake rates of metabolites are determined by kinetic rate laws and product secretion rates are predicted by the metabolic model. To compute fluxes using the metabolic model, COSMIC-dFBA first determines the number of metabolic states by inspecting uptake and production fluxes between various sampling intervals. Using these data, we then compute the fraction of cells in each phase, which provides a measure of state shift. We then identify the metabolites that show a significant difference in concentration between the identified states and train the cell state distribution predictor, a statistical model to predict state shift based on the prevailing bioreactor conditions. Using uptake and secretion rates inferred from spent media analysis, we then generate a priority list for metabolic tasks to determine the order of resource allocation of various cellular objectives for each identified state. A parameterized kinetic rate law is used to constrain nutrient uptake in each identified state. This information is then used to solve the metabolic model and predict the net uptake and secretion rates of all tracked metabolites. This framework accurately predicts concentration profiles and antibody titers in a diverse range of bioreactor conditions including glucose, amino acid, and oxygen depleted media. Therefore, this framework is a valuable resource for bioprocess characterization and optimization.

**Figure 1A:**
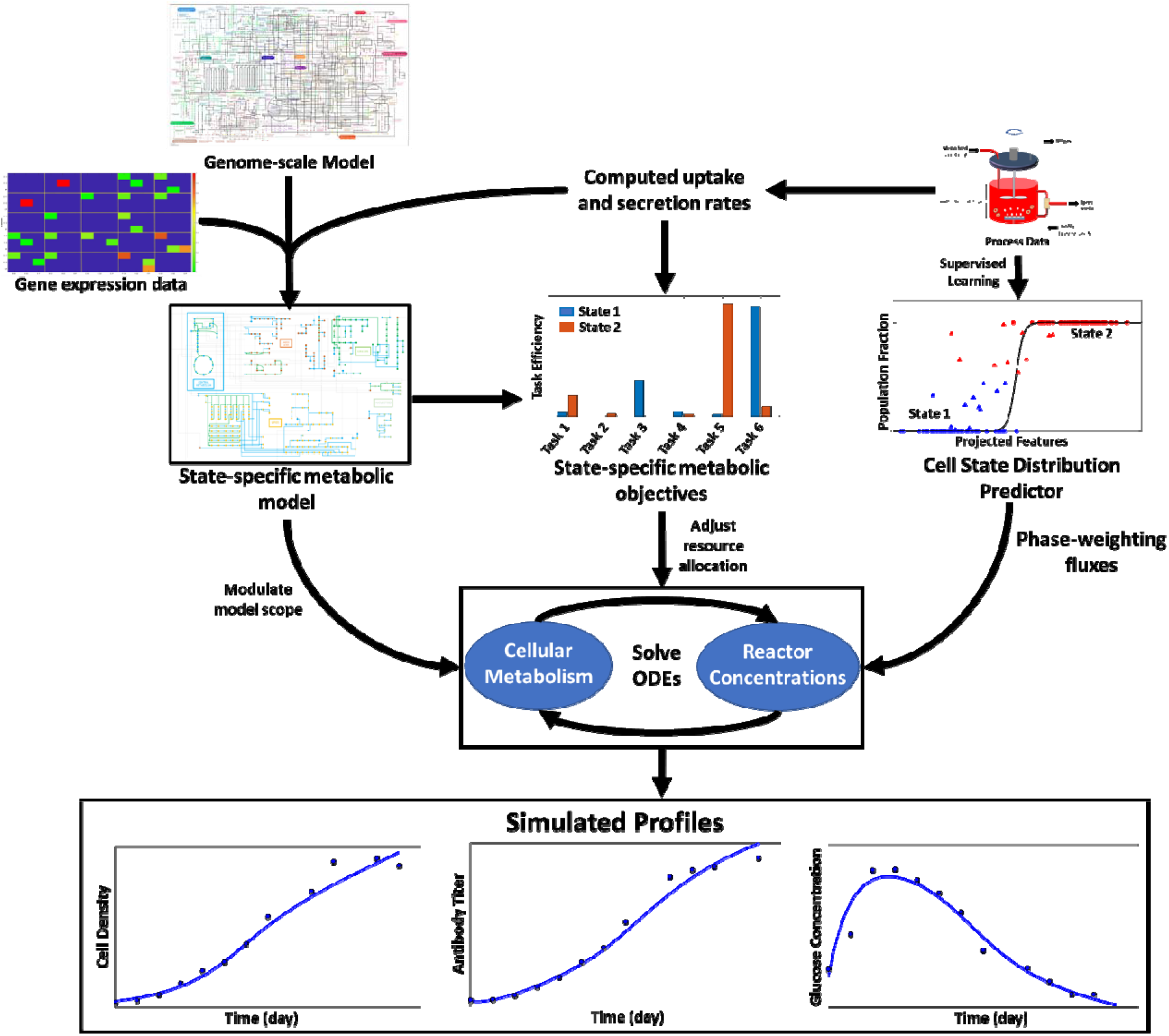
Overall workflow showing the pre-requisites and simulation approach used by COSMIC-dFBA. COSMIC-dFBA predicts metabolite concentration, cell density, and antibody titer profiles by solving a system of ordinary differential equations in which the rate of metabolite uptake/secretion is determined using a metabolic model. In order to accomplish this, three inputs must be specified. The first input is the state-specific metabolic model, which is derived from a genome-scale metabolic model by overlaying different types of -omics data (metabolomic, transcriptomic, or fluxomic data). The second requirement is the knowledge of state-specific cellular objectives encoding the allocation of nutrients into various products, which is inferred from metabolite uptake and secretion rates computed using spent media analysis. The third requirement is a cell state distribution predictor, a machine learning model that predicts the cell state based on prevailing conditions to adjust nutrient uptake by the metabolic model.

**Figure 1B:**
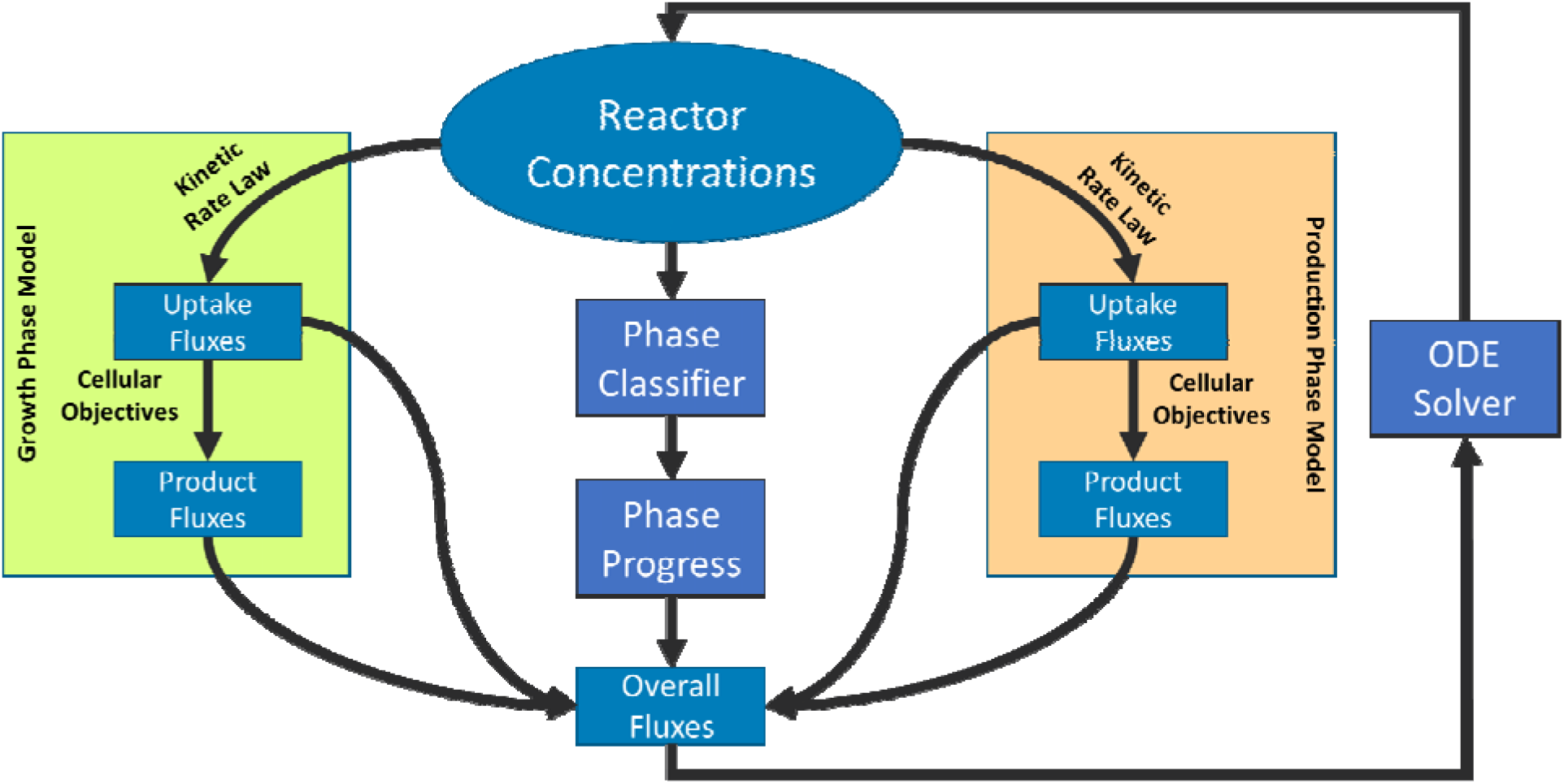
Computing instantaneous metabolic fluxes in COSMIC-dFBA. The system of ODE solved to update reactor metabolite concentrations requires uptake and secretion rates that are computed as a weighted average of metabolism from all possible metabolic states (growth and production states, in this case). The weights for the contributions are computed using the cell state distribution predictor. The fluxes corresponding to each metabolic state are solved by solving a multi-level flux balance analysis problem using the state-specific metabolic model, provided state-specific uptake rates (determined by reactor metabolite concentrations using a Monod-like equation), and specified cellular objectives. The net result is a set of flux distributions corresponding to various metabolic states. These flux distributions are averaged based on weights computed by the cell state distribution predictor to obtain the net uptake and secretion rates.

## 2. Results

### 2.1. The COSMIC-dFBA framework

**<u>C</u>**ellular **<u>O</u>**bjectives and **<u>S</u>**tate **<u>M</u>**odulation **<u>I</u>**n biorea**<u>C</u>**tors (COSMIC-dFBA) is a multi-scale hybrid dynamic flux balance analysis framework that predicts total cell density, antibody titer, and metabolite concentration profiles throughout a bioprocess. Figure 1 shows the schematic representation of COSMIC-dFBA along with the pre-requisites and dynamic inputs required for execution. We define a metabolic state (hereafter referred to as “state”) as the aggregate of nutrient uptake, afforded pathways for metabolism, and flux distribution into various products. The conceptual advancement by COSMIC-dFBA is the seamless transition between states in a dynamic bioprocess without the need for condition-specific parametrization of state transition. Because FBA is only applicable at metabolic steady-state, intracellular flux distributions are constrained via nutrient uptake rates in traditional dFBA. COSMIC-dFBA overcomes this limitation by assuming that overall metabolism in the reactor is a weighted average of metabolism of cells in various states. The cell state at any time point is predicted by the Cell State Distribution Predictor model based on instantaneous bioreactor conditions and feature metabolite concentrations using a supervised machine-learning classifier (See Methods section 4.4. The four prerequisites for executing COSMIC-dFBA include (i) state-specific metabolic models that contain limits on nutrient uptake and pathways available for metabolism, (ii) state-specific uptake kinetics that reflect the effects of changing gene expression on nutrient uptake in different cell states, (iii) state-specific metabolic tasks that encode the resource allocation in each cell state, and (iv) a machine learning model to predict population distribution among cell states based on the prevailing conditions in the bioreactor. The procedure for preparing these prerequisites is described in the Supplementary Methods.

COSMIC-dFBA simulates bioreactor metabolite and product concentrations by solving a system of ODEs describing the feeding, removal, and metabolism of nutrients and products in the bioreactor (Figure 1B). At each time point, the uptake rates and secretion rates are computed in three steps. First, uptake rates for all nutrients are calculated using computed kinetic rate laws for each state. Next, the computed nutrient uptake rates are used to constrain the respective state-specific metabolic models. The state-specific secretion rates are computed by solving the state-specific metabolic model using a multi-objective FBA. Finally, the average uptake and secretion rates are computed by weighting the computed state-specific uptake and secretion rates by the fraction of cells in each state, predicted by the cell state distribution predictor model based on prevalent bioreactor condition. These overall rates are then used to update the nutrient concentrations in the bioreactor.

### 2.2. Cellular objectives are cell state-specific

A PCA of computed fluxes revealed two distinct cellular metabolic states (state 1 and state 2) representing metabolism before day 3 and after day 10. We analyzed the computed state-specific uptake and secretion rates (see Methods section 4.3 and Supplementary methods section 1) in the context of the *i*CHO1766 metabolic model to quantify the changes in resource allocation associated with state shift. We first computed the task efficiencies (defined as the ratio of measured flux to maximum flux predicted by the metabolic model) for each secreted product and assigned priorities to each metabolic task (see Supplementary Methods). Figure 2 shows the task efficiency averaged across all reactor conditions for all measured metabolic byproducts in both states. We found that biomass formation and lactate secretion were the top two metabolic tasks in state 1, accounting for 88% of the consumed carbon and 40% of the consumed nitrogen, as quantified by FBA. Based on this, we call state 1 the “growth state”. The primary metabolic task in the production phase was antibody production, accounting for 73% of the consumed nitrogen. Based on this, we call state 2 the “production state”. Although the total cell density did not change for cells in the production state, the cell size steadily increased, suggesting that biomass precursors were being synthesized and accumulated. These findings demonstrate that metabolism qualitatively changes upon state shift in the bioprocess and motivates the need to incorporate approaches to account for cell state shifts and changing metabolic objectives in a dFBA simulation.

**Figure 2:**
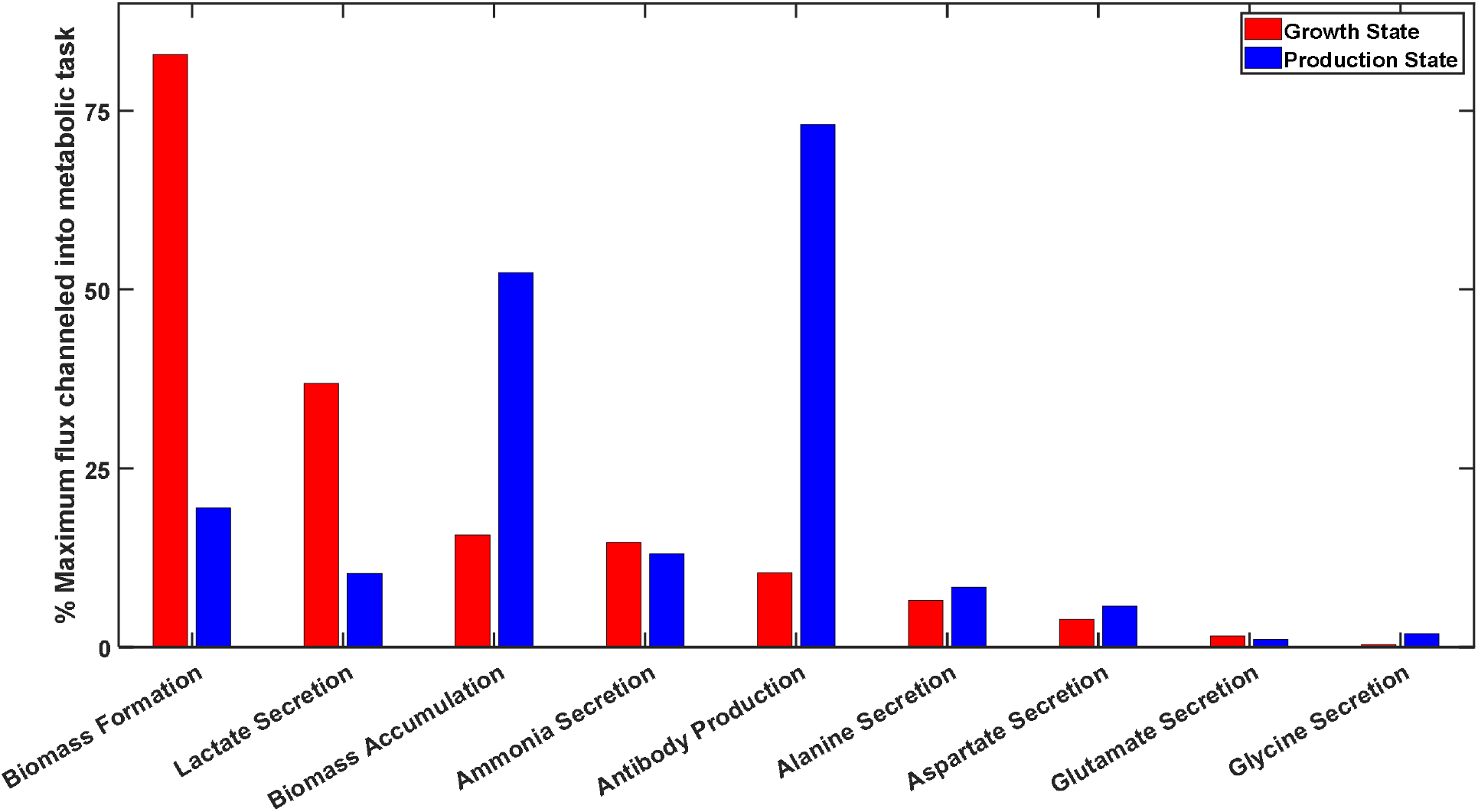
Resource allocation towards various metabolic tasks in the growth and production phases. Cells were predominantly in the growth state before day 3 and transitioned to the production state between day 3 and day 10. Past day 10, cells were primarily in the production state. Most of the cellular resources were channeled into biomass formation in the growth state and towards antibody production in the production state. Lactate was produced from glucose via glycolysis and from asparagine and glutamine via the anaplerotic pathways. Additional carbons were channeled into synthesizing biomass precursors in the production state, which were accumulated intracellularly. A similar fraction of consumed nitrogen was channeled into ammonia generation (via glutaminolysis and asparagine degradation) and alanine production via transamination in both states. Glycine production was significantly reduced in the production state.

### 2.3. The cell state distribution predictor captures phase-shifts driven by nutrient and oxygen depletion

To ensure that cell state is properly predicted by changes in reactor conditions when simulating a bioprocess, we developed a state classification model and trained it through a three-step workflow (Figure 3). The first step is to identify metabolites whose depletion correlates with the observed state shift. To accomplish this, we label each data measurement as either growth state, production state, or mixed state based on the state progression parameter computed concurrently with uptake and secretion rates (see Supplementary Methods section 1). The state progression parameter, p, represents the distribution of cell populations in each metabolic state with p = 0 indicating that all cells are in the growth state and p = 1 indicating that all cells are in the production state. Metabolite concentrations at time points with p < 0.2 (less than 20% of the cells in the production state) were considered to represent the growth state and at time points with p > 0.8 (more than 80% of the cells in the production state) were considered to represent the production state. We excluded metabolites whose media concentration increased over time as metabolic byproducts were constantly cleared from the bioreactor by perfusion and retain only those metabolites whose media concentration decreases by at least 50%. For the cell line and process considered here, the full list of features includes glucose, asparagine, and glutamine as the potential metabolite candidates in addition to oxygen level and bioreactor temperature (Supplementary Figure S1).

**Figure 3:**
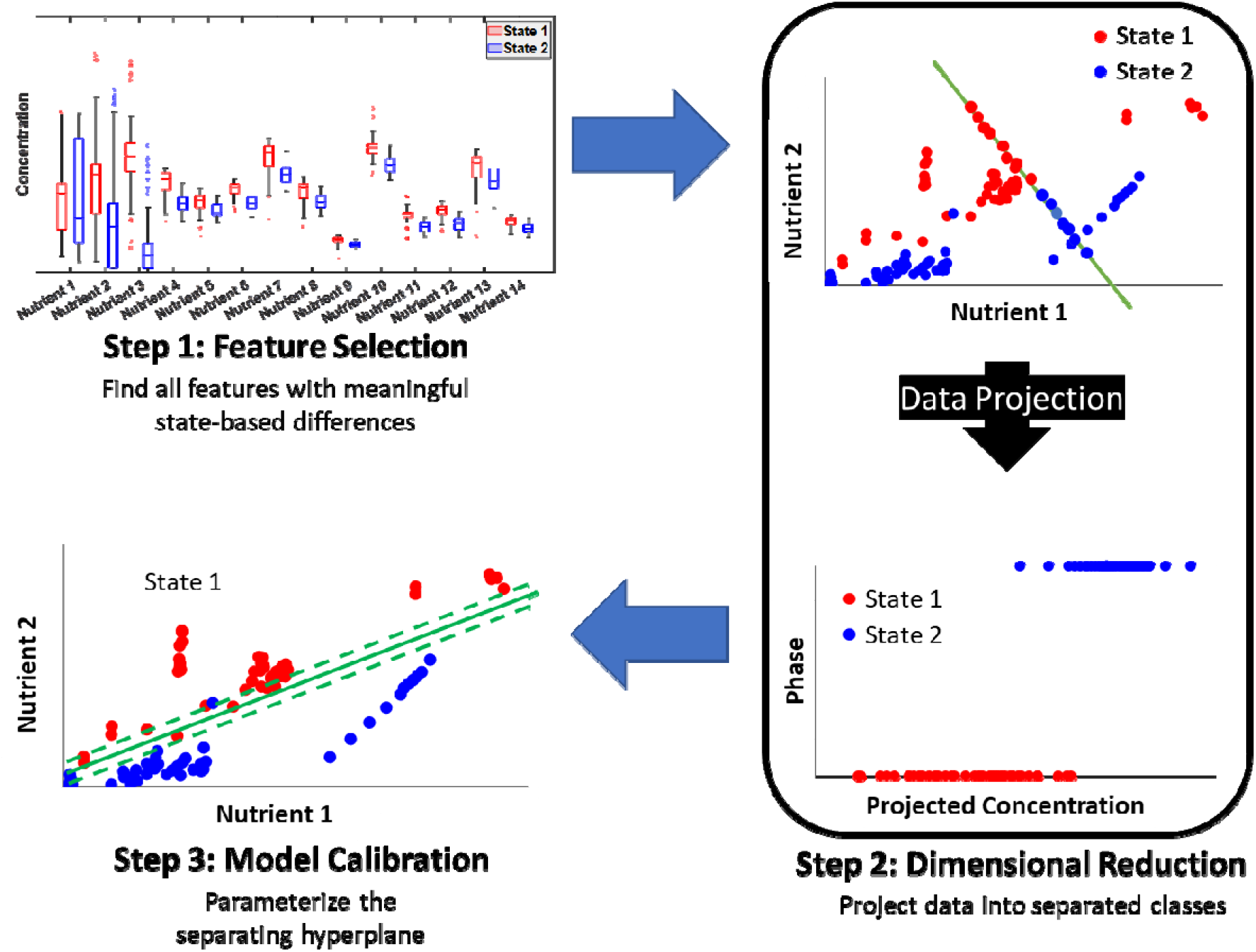
Training the phase classifier model to predict cell state based on bioreactor conditions

The second step in developing the phase classifier model is dimensional reduction with Linear Discriminant Analysis (LDA) to project the features to a lower dimensional space, such that projected features are correctly classified into the growth and production state. For this, we consider features corresponding to growth state and production states (p < 0.2 or p > 0.8) and ignore features corresponding to the mixed state. In the final step, we fit a logistic curve to model the relationship between projected features from all three states and the predicted state progression parameter. The resulting machine learning model intakes the prevailing bioreactor features and predicts cell state to determine the net uptake and secretion rates in the reactor (See Figure 1B).

The cell state distribution predictor model correctly predicted the state for 94 of 130 time points across all bioreactor growth conditions with an accuracy of 0.1 (difference between predicted and computed state is less than 0.1) and 118 of 130 time points with an accuracy of 0.2 (Figure 4). The model had a specificity of 0.78 and a sensitivity of 0.681. The F1-score was 0.731 and Matthews’ correlation coefficient was 0.454. In contrast, models based on a random classifier (cell state distribution assumed to be a random number between 0 and 1) had a Matthews’ correlation coefficient of -0.72, indicating that the state prediction by the trained model significantly outperformed random chance (permutation test, p-value < 10^-6^). The model correctly identified state shifts associated with the depletion of asparagine, glutamine, and glucose, even in growth conditions with altered amino acid and glucose availability. In cases with altered oxygen availability, the model correctly identified the cell state for all data except those between days 6 to 8. This was because the cells had already transitioned into the production state in the oxygen-depleted condition before the feature metabolites were sufficiently depleted for the model to identify and predict a state shift, leading to a false negative prediction. Other cells had not transitioned to the production state despite depletion of the feature metabolites in the high-oxygen condition, leading to a false positive prediction by the phase classifier. Except for a small number of extreme conditions, the model robustly predicts cell state shifts arising from nutrient depletion within the bioreactor across a wide range of conditions.

**Figure 4:**
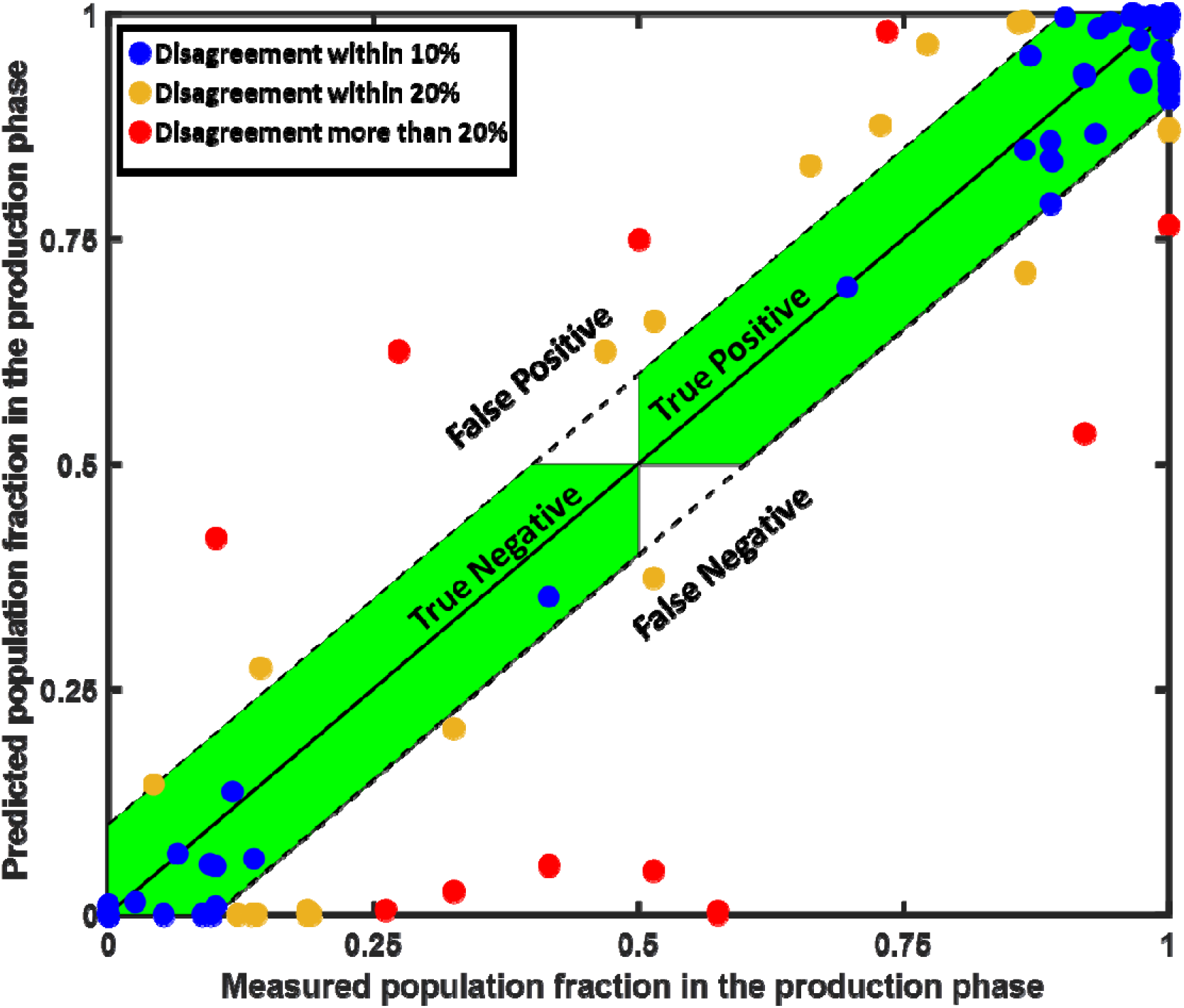
Comparison of model-predicted and measured population fractions in the production phase. The Blue dots represent the data points that were correctly identified to be in either the growth or production phase with a 10% margin of error. The orange dots represent the correctly predicted phases with a 20% margin of error. The red dots represent data that were incorrectly predicted by the phase classifier model.

**Figure 5A:**
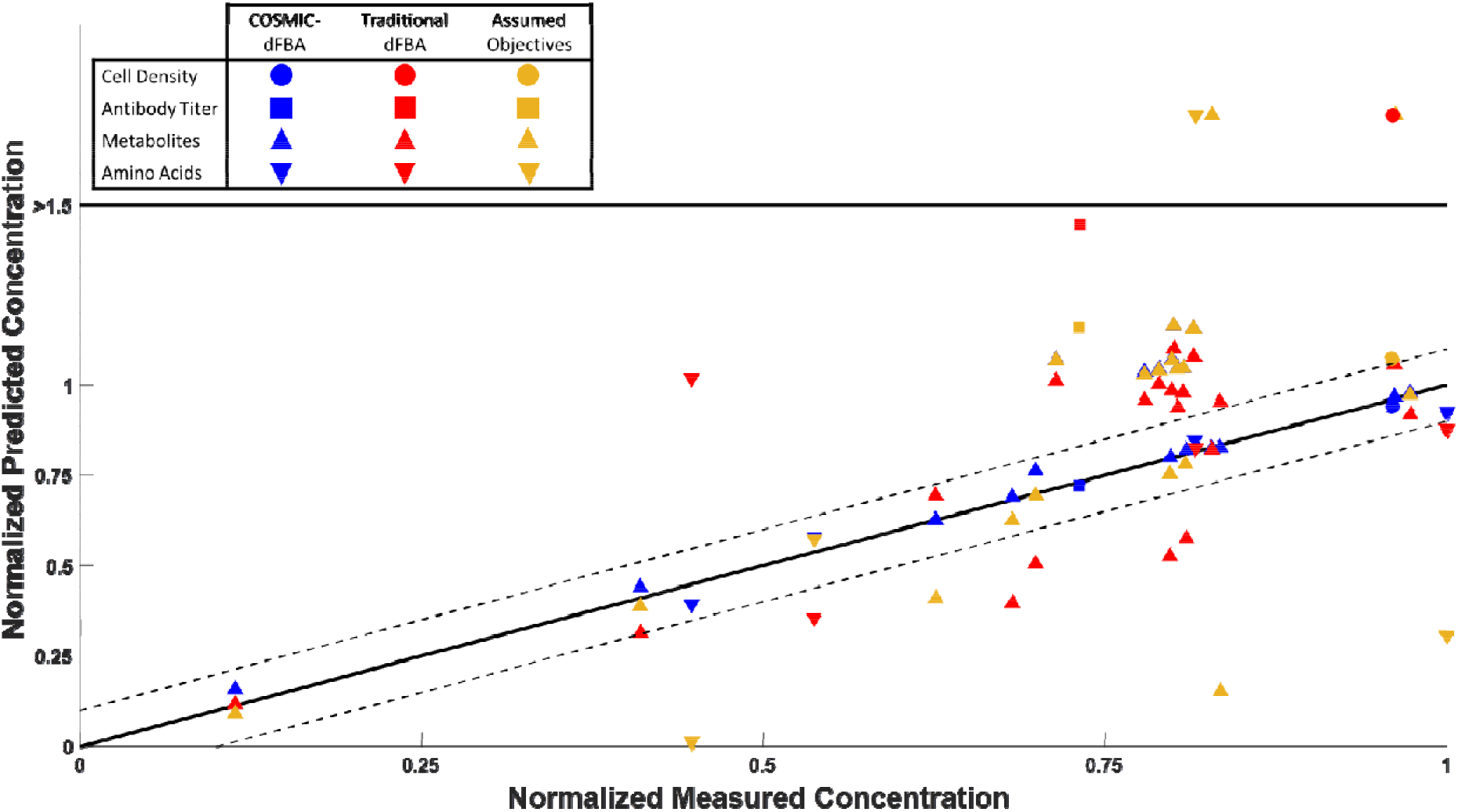
Consistency of measured and predicted concentrations on day 13 for amino acid (downward triangles), glycolytic metabolites (upward triangles), cell density (circles) and antibody titer (square) using COSMIC-dFBA (blue markers), a standard dFBA algorithm with specified cellular objectives and phase switch at a fixed time point (red markers), and a standard dFBA algorithm with the phase classifier from COSMIC-dFBA but assuming maximize biomass objective during the growth phase and maximize antibody production objective in the production phase (orange markers).

**Figure 5B:**
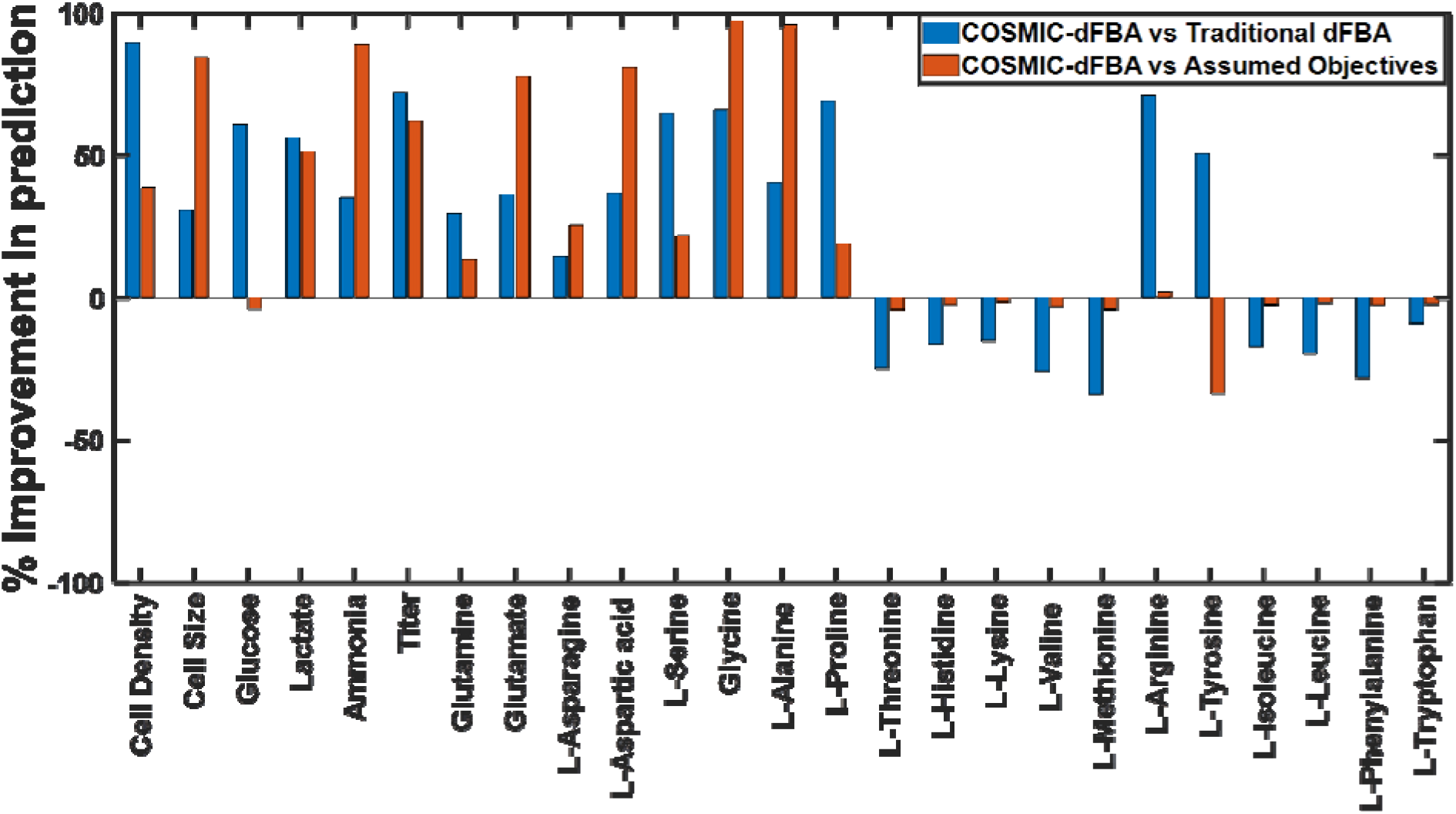
Improvement in predictions by COSMIC-dFBA compared to models with no classifier or assumed objective functions.

### 2.4. The dFBA algorithm accurately predicts concentration profiles

We simulated the cell density, glucose, lactate, antibody, and 17 amino acid concentration profiles over the 13-day perfusion bioprocess run across 8 different media conditions (Supplementary Figures S2 – S11). To evaluate its performance, we compared the concentration profiles, predicted using COSMIC-dFBA with two implementations of traditional dFBA. In th first case (referred to as the “traditional dFBA case”), we retained the cellular objectives, but assumed that phase transition coincided with the hypothermic shift. In the second case (referred to as the “assumed objective case”), we retained the phase classifier from COSMIC-dFBA, but assumed that the cells only maximize biomass during the growth phase and maximize antibody production in the production phase. Figure 4A compares the day 13 concentration predictions by COSMIC-dFBA and the two test cases. We found that COSMIC-dFBA significantly outperformed both test cases based on traditional dFBA, thus highlighting the need to account for changing bioreactor conditions and metabolic tasks. We also evaluated the improvement in prediction (defined as the mean fractional reduction in disagreement between predicted and measured concentrations over the course of the bioprocess) as a measure of how well the concentration profiles predicted by each algorithm agree with the experimental data. From this we found that the concentration profiles predicted by COSMIC-dFBA for cell density, antibody titer, glucose, lactate, glutamine, and glutamate were in better agreement with the measured data than the traditional dFBA cases (Figure 4B). However, the standard dFBA test cases better predicted the consumption of several essential amino acids.

The traditional dFBA case greatly overestimated the final cell density in all eight growth conditions. This was because the traditional dFBA case assumed that the entire cell population transitioned from the growth phase to the production phase when the hypothermic shift was applied, regardless of the bioreactor conditions. Thus, this case failed to account for the redistribution of metabolic fluxes and a shift from cell growth to antibody production when key metabolites were depleted early, particularly in the low glucose and low amino acid cases. This led to an extended growth phase in all eight conditions, and a higher cell density at the end of the growth phase. Consequently, this approach predicted a higher antibody titer in all growth conditions. On the other hand, the final cell density predicted using the “assumed objectives” case was only 14.8% higher than those predicted using COSMIC-dFBA. This agreement between COSMIC-dFBA and the “assumed objectives” case arises from the fact that the assumed maximization of biomass formation is close to the actual metabolism of the cells, which channels, on average, 82% of the resources towards biomass production in the growth phase (Figure 2). This implementation also assumed that all available resources were channeled into antibody production in the production phase, whereas the experimental data suggests that 25% of the resources were channeled into other cellular processes. This led to a dramatic overprediction of antibody titer in the bioreactor. Overall, these comparisons demonstrate the importance of the two integral components of COSMIC-dFBA (the phase classifier and comprehensive accounting of metabolic tasks), which contribute to the algorithm’s superior predictive capabilities compared to existing dFBA-based bioprocess modeling frameworks.

## 3. Discussion

This study presents COSMIC-dFBA, a multi-scale dynamic flux balance analysis framework that combines machine learning and mechanistic modeling techniques to simulate cell behavior in a perfusion bioprocess and predict metabolic shifts in response to changing bioreactor conditions. This framework operates at two scales: the bioreactor scale and the cellular scale. The cellular scale interfaces with the bioreactor scale using the cell state distribution predictor that determines the distributions of cell populations in various states based on prevailing bioreactor conditions. Based on the determined cell state, nutrients are consumed according to previously parameterized kinetic rate laws, and consumed nutrients are channeled into appropriate state-specific metabolic tasks (e.g., cell growth, antibody production, etc.). This yields the net instantaneous production and consumption rates of all metabolites in the bioreactor, which are then used to update the bioreactor concentrations by solving a system of ODEs. Leveraging the metabolic model provides a mechanistic relationship between nutrient uptake and product secretion as well as additional pathways through which metabolic flux is diverted to generate byproducts. By dynamically adjusting product yields, this framework always ensures that nutrient consumption and product formation in the bioreactor satisfy conservation of mass and are thermodynamically feasible, which is not always the case when modeling a bioprocess using empirical models. Unlike previous dFBA approaches (Nolan and Lee, 2011), COSMIC-dFBA does not need to solve any quadratic programming problems, which considerably decreases the computational cost. This permits the use of genome-scale metabolic models for dFBA, which increases generalizability. Incorporating the means to modulate cellular resource allocation using a hybrid modeling paradigm improves fidelity without the need for developing detailed mechanistic models such as whole-cell models or ME-models. Furthermore, by using an adaptive time step, a desired integration accuracy can be ensured without resorting to collocation (St John et al., 2017), which significantly reduces the number of time-steps and by extension, the number of times the FBA problem must be solved (de Oliveira et al., 2023; Zhuang et al., 2011).

COSMIC-dFBA is particularly versatile in that it only requires the usual data typically collected during a bioprocess to train the model. Uptake and secretion rates were computed from metabolite concentration profiles and analyzed to determine phase-specific resource allocation to identify the major metabolic tasks prioritized by the cell in various states, whereas phase shifts were predicted based on reactor metabolite concentrations and temperature shifts. Other types of omics data can be readily incorporated to minimize manual interventions. For example, integrating gene expression data enables extraction of context-specific metabolic models (Gustafsson et al., 2023; Opdam et al., 2017), which have been previously shown to vary between process phases. Transcriptomics data can also suggest metabolic tasks not captured by exo-metabolomics data (Helen et al., 2022; Masson et al., 2023; Richelle et al., 2021). Proteomics data can be incorporated to correlate changes in transporter abundance with phase shifts (Colijn et al., 2009; Sanchez et al., 2017; Tian and Reed, 2018; Yeo et al., 2020), which modulates the maximum uptake rate of nutrients in each process phase.

The key strength of COSMIC-dFBA is the ability to learn from additional experimental data that allows it to predict newer states. Our analyses indicate that the cell state distribution predictor is a vital component of this framework that smoothly modulates state shifts using a single layer perceptron (linear combination of inputs combined with a logistic activation function). The choice of activation function was based on previous efforts to model cellular signal transduction (Samaga and Klamt, 2013; Wynn et al., 2012) and gene activation (Ay and Arnosti, 2011) in response to changing environmental conditions within the bioreactor. The main drawback of this approach is that the framework cannot automatically determine the cause of the state shift (arising from nutrient depletion, temperature shift, oxygen limitation, etc.) and assumes that all phase shifts are of the same nature. In the current implementation of COSMIC-dFBA, we circumvent this by defining the cellular objectives for each type of phase shift in advance. However, automated prediction of changes in metabolic task priorities in response to phase shifts will require an overlay of the signaling (Lin et al., 2022; Sompairac et al., 2019) and gene expression networks (Pio et al., 2022) on to existing models of metabolism in the absence of fully descriptive whole-cell models (Ahn-Horst et al., 2022; Karr et al., 2012). Such models will expand the predictive capabilities of COSMIC-dFBA to predict heterogeneity in cell populations in large-scale bioreactors arising from non-homogeneous mixing and poor local oxygen transfer. That will allow the framework to predict and correct the potential detriments to process yield and productivity upon scale-up to manufacturing scales. Despite these limitations, COSMIC-dFBA significantly outperforms traditional dFBA in its current form. The ability to model dynamic metabolism uniquely positions this framework for applications in bioprocesses with metabolic shifts.

## 4. Methods

### 4.1. Cell culture and process data acquisition

A stable, clonally derived Chinese hamster ovary (CHO) cell line expressing a non-glycosylated recombinant protein was thawed and scaled up in proprietary growth media to generate sufficient cell mass to inoculate a production perfusion bioreactor. The production bioreactors were operated in 3 L stirred tank bioreactors with a 1.5 L working volume for 13 days using proprietary chemically defined media. Bioreactors were inoculated in the same basal production media. Perfusion was performed using alternating tangential flow filtration starting at Day 0 at a perfusion rate of 1 bioreactor volume per day for a duration of 13 days. On Day 8, the temperature setpoint was decreased for the remaining duration of the experiment. The experimental conditions were set up following a Box Behnken DOE varying dissolved oxygen, perfusion media amino acid levels, and perfusion media glucose concentration as shown in Supplementary Table ST1.

Bioreactor parameters, such as agitation, dissolved oxygen concentration, pH, and temperature were monitored and controlled through a DeltaV controller (Emerson, St. Louis, MO, USA). The pH was controlled through CO_2_ or 1 M Na_2_CO_3_ addition. Dissolved oxygen was maintained by sparging oxygen through a drilled pipe and a sintered sparger. Additionally, inline off-gas O_2_ and CO_2_ were monitored using the BlueSens BlueVary gas sensor (BlueSens, Wood Dale, IL, USA). The daily sampling procedure consisted of cell density and viability using a Cedex HiRes analyzer (Roche Diagnostics, Indianapolis, IN, USA), metabolites (lactate, glucose, glutamine, glutamate, and ammonium) from a Cedex Bio HT analyzer (Roche Diagnostics, Indianapolis, IN, USA), osmolality using the Advanced Instruments OsmoPRO (Advanced Instruments, Norwood, MA, USA), and external pH, pCO_2_, and pO_2_ using a Siemens RAPIDLab 1260 (Siemens Healthineers, Erlangen, Germany). Daily clarified samples for each reactor were analyzed for titer via HPLC. Amino acid concentrations were determined as follows: cell culture supernatant samples were filtered through a 0.2µm filter then diluted properly with 18 mM HCl and mixed with the internal standard mixture containing heavy isotope labeled amino acids. An UHPLC system Agilent 1290 (Agilent Technologies, Santa Clara, CA, USA) equipped with a reversed phase C18 column (Agilent Poroshell 120 SB-C18, 1.9 µm, 2.1 mm × 100 mm) was used for components separation. The mobile phases used were water (A) and acetonitrile (B) in 0.2% heptafluorobutyric acid (HFBA). Targeted quantitation data were acquired using the dynamic Multiple Reaction Monitoring (MRM) mode on an Agilent 6490 Triple Quadrupole mass spectrometer. Agilent MassHunter B.08.00 was used for data acquisition and data analysis.

### 4.2. Metabolic model and data processing

iCHO1766 was used as the base metabolic model (Hefzi et al., 2016). The protein secretory pathway (Gutierrez et al., 2020) was appended to iCHO1766 to accurately model the precursor and energy demands for antibody synthesis and secretion. Two phases were identified using the concentration data. The growth rate, antibody specific productivity, uptake and secretion rates of all measured metabolites, and the fraction of cell population in each phase were computed from the concentration profiles using nonlinear regression as described in the supplementary methods. The computed fluxes in each growth condition are reported in Supplementary Table ST3.

### 4.3. Inferring state-specific metabolic task objectives and priorities

State-specific metabolic flux distributions were modulated in terms of metabolic tasks and task efficiencies. Each state-specific model was calibrated as described in the supplementary material. Briefly, all measured quantities were classified into either nutrients (consumed by cells) or byproducts (generated by cells) in each phase. All secreted byproducts were considered “metabolic tasks” and their priority order was determined in an iterative manner. First, the uptake rates of nutrients were fixed in the metabolic model. Following this, the flux through each metabolic task was individually maximized using Flux Balance Analysis (FBA) (Varma and Palsson, 1994). Task efficiency for each metabolic task was computed as the ratio of measured flux through the metabolic task to the maximum flux predicted using FBA. The task with the highest efficiency was considered the highest priority task as it reflects the maximal nutrient utilization towards this task and its corresponding efficiency was stored. To find the next priority task, the experimentally measured flux through the previous task was enforced as a lower bound in the metabolic model and that task was removed from the list of metabolic tasks to be evaluated. Following this, the task efficiency calculation steps were repeated to identify the next highest priority task. This loop was repeated until all metabolic tasks were ordered. The list of state-specific metabolic tasks and their corresponding task efficiencies are reported in Supplementary Table ST4.

### 4.4. Training the cell state distribution predictor

The cell state distribution predictor is a machine-learning model that predicts cell state based on bioreactor conditions. Using bioreactor media concentrations, partial pressure of oxygen, and the temperature of the bioreactor as inputs, the phase classifier model is trained using the three-step process depicted in Figure 3 and predicts the fraction of cell population in the production state. For each condition considered in this study, time points were classified into either growth state, production state, or mixed populations based on whether the fraction of cells in the production state were less than 20%, greater than 80%, or somewhere in between, respectively. Concentrations of all metabolites were grouped into these three classes and plotted to identify metabolites correlated with phase shifts. Candidate metabolites were chosen such that their (a) median concentrations changed drastically between the growth and production states, and (b) they were depleted, or close to depleted in the production state. Oxygen and temperature were included to account for premature state shifts arising from hypoxic (Zeh et al., 2021) and hypothermic shifts (Wulhfard et al., 2008). The second step is to reduce the dimensionality of the data such that the growth and production state data are separated into distinct clusters. We used Linear Discriminant Analysis (LDA) to achieve this. The projected concentration *w* is related to feature *i* (metabolite concentration, partial pressure of oxygen, or temperature) via a weighted linear combination using weights *k_i_* using Equation (1):

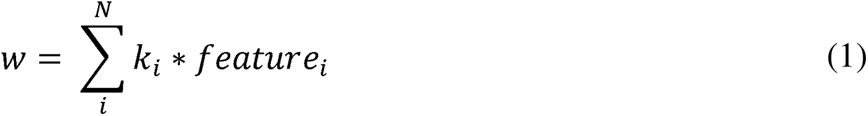

Following this, projected concentrations were computed for all three classes and logistic regression was performed to compute the parameters *a* and *h*, representing the steepness of the transition and the bias, respectively and model the transition from growth to production state using Equation (2):

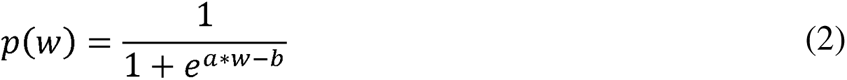

### 4.5. Simulating metabolite concentration profiles and culture parameters using COSMIC-dFBA

COSMIC-dFBA simulates bioreactor concentration profiles by solving the following initial value problem (IVP) for cell density (*X*(*t*)), cell size (*S*(*t*)), and concentration of metabolite *i* (*C_i_*(*t*)) from time *t*_0_ to *t*_f_:

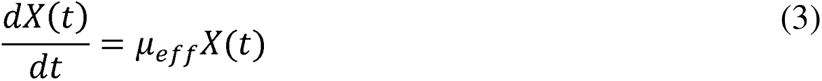

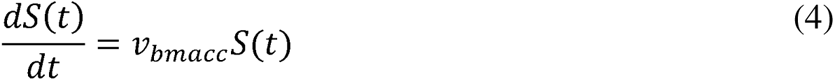

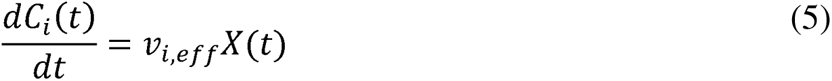

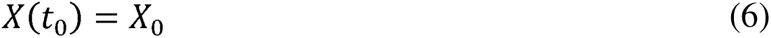

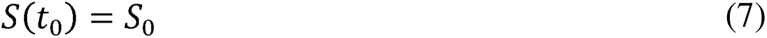

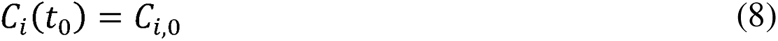

In Equations (3), (4), and (5), µ*_eff_*, *v_bmacc,eff_*, and *v_i,eff_* represent the effective growth rate, effective biomass accumulation rate, and the effective uptake/secretion rate of metabolite *i*, and are related to the growth and production phase fluxes via the population fraction parameter *p*(*t*) computed using Equations (1) and (2):

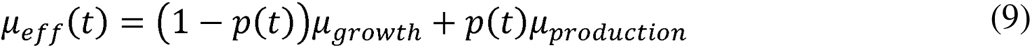

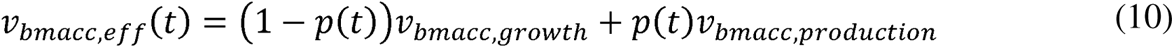

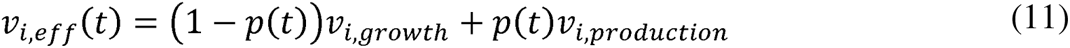

The above IVP is solved using the Bulirsch-Stoer algorithm (Bulirsch and Stoer, 1966) with adaptive step-size control (Deuflhard, 1983) to reduce the number of times the metabolic model must be solved without loss of accuracy. COSMIC-dFBA is encoded and executed in MATLAB^TM^. The source code is provided as a zip file in the supplementary material.

## Acknowledgements

This work was supported by a generous grant from Amgen.

