## Supplementary Material for "COSMIC-dFBA: A novel multi-scale hybrid framework for bioprocess modeling"

**Supplementary Methods**

**Flux and Phase Calculation**

Fluxes (growth rate, uptake, and secretion rates) associated with all quantities (cell density, cell size, antibody, metabolites, and amino acids) were computed using nonlinear regression so as to minimize the variance-weighted sum of squared deviation of predicted quantities from their experimentally measured values. Metabolite concentrations are predicted using a two-state phenotypic model. Cells are assumed to be either in the growth phase or in the stationary phase, each represented by a unique set of fluxes. We define $v_{i}^{\left( k \right)}$ as the fluxes associated with quantity $i$ in phase $k$ (either $(growth)$ or $(stat)$). The objective is to identify the set of $v_{i}^{\left( k \right)}$ that recapitulates the measured concentrations of all quantities at all sampling times ($t_{0}, t_{1},\ldots,t_{M}$). We first divide the process into M intervals based on sampling times such that interval $m$ represents the time interval $[t_{m-1},t_{m}]$. For each time interval, we define a parameter $0\leq p_{m}\leq1$ that represents the fraction of the cell population in the stationary phase. Thus, the net flux in interval $m$ associated with quantity $i$ is computed using Equation (1).

$$v_{i,m}=\left( 1-p_{m} \right)v_{i}^{\left( growth \right)}+p_{m}v_{i}^{\left( stat \right)}$$

(1)

The time-course profile of all quantities in the interval $m$ are then computed by solving the system of ODEs described in Equation (2).

$$\frac{dC_{i}}{dt}=F\left( C_{i}^{\left( in \right)}-\eta C_{i} \right)+v_{i,m}C_{1}$$

(2)

In the above Equation, $C_{1}$ refers to the cell density in the reactor, $C_{i}^{\left( in \right)}$ denotes the concentration of quantity $i$ in the perfusion medium, $F$ denotes the perfusion rate (equal to 1 $d^{-1}$), and $\eta$ is the removal fraction of the quantities from the bioreactor. $\eta$ equals 0 for cell density and cell size at all time points, and for antibody before day 8. In all other cases, $\eta$ equals 1.

Thus, the parameters $v_{i}^{\left( k \right)}$ and $p_{m}$ are computed by solving the following nonlinear optimization problem:

|  | $\min_{v_{i}^{\left( k \right)},p_{m}} \sum_{i=1}^{N} \sum_{m=1}^{M} \left( \frac{C_{i}\left( t_{m} \right)-C_{i}^{\left( meas \right)}\left( t_{m} \right)}{\sigma_{i,m}} \right)^{2}$ |  |  |
| --- | --- | --- | --- |
| Subject to: | $\frac{dC_{i}}{dt}=F\left( C_{i}^{\left( in \right)}-\eta C_{i} \right)+v_{i,m}C_{1}$ | $\forall t\in[t_{m-1},t_{m}]$ $1\leq m\leq M$ | (2) |
|  | $v_{i,m}=\left( 1-p_{m} \right)v_{i}^{\left( growth \right)}+p_{m}v_{i}^{\left( stat \right)}$ | $1\leq m\leq M$ $1\leq i\leq N$ | (1) |
|  | $0\leq p_{m}\leq1$ | $1\leq m\leq M$ | (3) |
|  | $p_{m}\geq p_{m-1}$ | $1\leq m\leq M$ | (4) |
|  | $p_{1}=0$ |  | (5) |
|  | $p_{M}=1$ |  | (6) |
|  | $C_{i}\left( t_{0} \right)=C_{0,i}$ | $1\leq i\leq N$ | (7) |
|  | $C_{i}\left( t \right)\geq0$ | $1\leq i\leq N$ | (8) |
|  | $v_{i}^{\left( k \right)}\mathbb{\in R}$ | $1\leq i\leq N$ $k\in\{growth,stat\}$ | (9) |

Kinetic parameters need only be computed for fluxes corresponding to taken up quantities. The kinetic rate law is defined using a Michaelis-Menten-type equation. We fix the parameters $p_{m}$ and all $v_{i}^{\left( k \right)}\geq0$, replace all $v_{i}^{\left( k \right)}<0$ with the kinetic rate law and solve the above optimization problem again to compute the $v_{MAX}$ and $K_{M}$ terms to modulate the uptake of quantities based on the reactor concentration. The NLP problem was solved using the fmincon function within the Optimization Toolbox in MATLAB^TM^.

**Mining metabolic tasks and task efficiencies**

We first classify the growth rate, antibody productivity, and the secreted metabolites in each phase as metabolic tasks. In order to accurately simulate the intracellular flux distribution for a given set of nutrient uptake rates, we must rank the metabolic tasks based on resource allocation determined using the *i*CHO1766 metabolic model. We also define “task efficiency” as the ratio of measured flux through a metabolic task to the maximum flux predicted using the metabolic model. The metabolic task priority and efficiencies for each phase are computed using the following algorithm.

| Step 1: | Impose the uptake rates of measured metabolites as bounds in the metabolic model. |
| --- | --- |
| Step 2: | Set $L = list of measured products$, $N = number of measured products$ and $i = 0$. |
| Step 3: | Set $i = i+1$ |
| Step 4: | Using FBA, compute the maximum flux through each product in $L$. |
| Step 5: | Compute $task efficiency =\frac{measured flux}{max flux}$ for each product in $L$. |
| Step 6: | Assign the product with the highest task efficiency to $Priority i$. Assign the corresponding task efficiency to $Efficiency i$. |
| Step 7: | Set the lower bound of flux through $Priority i$ to the measured flux. Remove Priority i from L. |
| Step 8: | If $L$ is empty, then STOP. Otherwise, repeat Steps 3 – 7. |
